# Rethinking Pleijel’s (1995) characters under a hierarchical point of view

**DOI:** 10.1101/2020.10.19.345330

**Authors:** Mathieu G. Faure-Brac, Valentin Rineau, René Zaragüeta Bagils

## Abstract

In 1995, Pleijel raised an issue on the coding of complex (multistate) characters: to find a unique coding approach satisfying both informational and semantical criteria. Following Pleijel’s work, this study aims to propose a new answer to this problematic. We proposed here to use hierarchical characters instead of classical partitional ones as hierarchy allows to deal effectively with the different uncertainties met by phylogeneticists since the beginning of phylogenetic data matrices: missing data, inapplicable data and polymorphism. We translated all previous proposed approaches into hierarchical ones and add three new approaches, only coded using hierarchy. Using phylogenetic and show with different metric how one of these new approaches displaces the other ones. The results from this study then supports the idea than phylogenetic characters should be coded using only hierarchies, as they offer a better management of uncertainties and propose new approaches more informative.

## Introducon

Since the publication of Phylogenetic Systematics (Hennig, 1966), many debates continue to animate the community of phylogeneticists on theoretical grounds. One of these most fundamental debates approaches the question of characters. Many authors questioned the nature of phylogenetic data (Pimentel & Riggins, 1987), their formalism (Sereno, 2007), and their representation (Cao et al., 2007; Mavrodiev et al., 2019; Prin, 2012; Williams & Ebach, 2006). Others tried to find the best practices to adopt about the coding of complex characters, especially questioning about the validity of multistate coding towards the absence/presence coding (Brazeau, 2011; Forey & Kitching, 2000; Hawkins et al., 1997; Lipscomb, 1992; Maddison, 1993; Platnick et al., 1991; Pleijel, 1995; Strong & Lipscomb, 1999).

These fruitful debates have led to the publication of numerous methodological studies, especially about coding approaches. Coding approaches are fundamental in systematic practices. They are the translation of observations of taxa into a symbolic representation suitable for an algorithmic process, the phylogenetic reconstruction. Coding characters being one of the first steps in phylogenetic reconstruction (Hennig, 1966), its execution impacts the entire rest of the analysis. Using one or another coding approach is, then, especially important to be carefully chosen.

Here, we focus on Pleijel’s (1995) study about issues in coding practices. In his paper, Pleijel compared various coding approaches of a complex morphological feature. In his fictional example, a specified morphological feature, named feature X, is either present or absent in taxa submitted to cladistic analysis. If it is present, it has one among four possible morphologies: black and round, striped and round, black and squared and striped and squared (fig 1). Pleijel proposed four different coding approaches in order to find the most suitable one to reconstruct phylogenies, using different criteria to compare them. He concluded by advocating for the coding approach which involves only the absence/presence (a/p hereinafter) of the different morphologies of the feature X, coded into binary characters, due to a better management of missing entries, as it avoids them, and of hierarchical linkage, as any new taxon can be added without implying any changes in the coding approach. However, this approach does not solve all issues raised by Pleijel.

Specifically, it violates the assumption of logical independency of characters(Vogt, 2018; Wilkinson, 1995). Characters are necessarily linked, as they are carried by the same organism. Hence, they necessarily co-evolve in a certain degree. This dependency is the biological one and is desired to recover phylogenetic relationships (Vogt, 2018; Wilkinson, 1995). However, the characters must not influence the way of coding the others because it will overweight their information. This dependency is the logical one and should be avoided (Vogt, 2018; Wilkinson, 1995). A full a/p approach violates this principle as each combination is a different variation of the same feature, to code them and the feature itself as so many different binary a/p characters violates this principle. This assumption is fundamental in cladistics, where the formalisation of each character hypothesis has to be independent of all others (de Pinna, 1991; Nelson, 1994; Pleijel, 1995; Prin, 2012). However, in the case of the a/p approach, there is a clear violation of this rule, as each variation of feature X relates to the feature itself and is a dependant state but coded as different characters (Zaragüeta Bagils & Bourdon, 2007). Moreover, this approach does not allow to code effectively hypotheses without loss of data (information retrieval *sensu* Pleijel (1995)). Each combination of shape and pigmentation of feature X is treated as a disconnected character from the others. There is no link between them and each one of them is judged without any reference to an alternative state. On his four criteria, the a/p coding approach fulfils only two of them. Pleijel does not succeed in his ambition to find the best coding approach and could only argue his favour for one among all, as all of his criteria are equally important. Five years later, Forey & Kitching (2000) proposed three new coding approaches to solve Pleijel’s issues, but their proposals did not solve the difficulties raised by Pleijel. The aim of this work is, then, to find a way to solve the issues arising from the coding of complex features raised by Pleijel.

**Figure 1.**
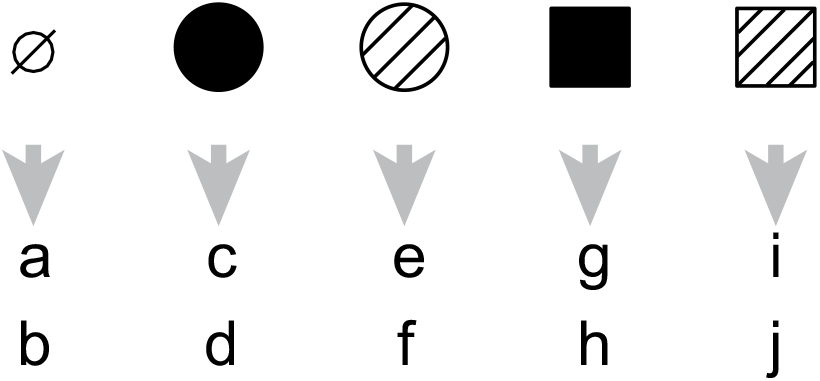
The different morphologies of feature X and the taxa bearing them. Taxa are named a to j. From left to right, the different morphologies are named as: absence of feature X, black rounded feature X, striped rounded feature X, black squared feature X and striped squared feature X.

An important step of the phylogenetic reconstruction, prior of the analysis itself, is then to translate the observations from taxa into a suitable form for a numerical analysis: characters in the data matrix. There is an underlying assumption in matrices, namely that the characters they store are systematically defined according to the particular type of mathematical set that is the partition (Colless, 1985; Farris, 1970; Pimentel & Riggins, 1987; Pogue & Mickevich, 1990). In set theory, a partition of a set is a set of subsets such that the intersection of any two of the subsets is empty and where every element is included in exactly one subset (Barthélémy & Guénoche, 1988). Partitions can be represented as matrices and as Euler’s diagrams. Our hypothesis is that the issues raised by Pleijel are caused by the use of partitions as structures for the representation of phylogenetic characters. Indeed, some authors have already shown that the conventional data matrix is incapable of effectively containing hypotheses of phylogenetic traits (Cao et al., 2007; Mavrodiev et al., 2019; Williams & Ebach, 2006). Moreover, Cao et al. (2007) identified the problem not as the data matrix itself, but as its partitional structure and proposed a model of matrix able to contain information in another kind of mathematical structure: hierarchies. It follows that none of the proposals of coding approaches from the literature are fully satisfactory because they are all thought in a partition framework.

Pleijel’s issues were always thought with the *a priori* that the characters must be conceived as partitions, as it is since Hennig (1966) and then in parsimony (Farris, 1977) or in Bayesian inference and Maximum likelihood (Felsenstein, 1973). It is interesting to note that there is another kind of mathematical set used in phylogenies. The very mathematical structure of any phylogenetic tree in cladistics is a hierarchy. A hierarchy is a set of subsets such that the intersection of any two of the subsets is empty or one of the two subsets, and where every element is included in at least one subset (Barthélémy & Guénoche, 1988). Hierarchies can be represented as particular matrices (Cao et al., 2007) and as Euler’s diagrams, but also as classificatory trees, such as cladograms. Characters are phylogenetic hypotheses, and carry within them the phylogenetic information that is then analysed to reconstruct a tree. One may agree that using partitions or hierarchies as mathematical structures in which are formalized the characters do not convey the same information content as they do not convey the same assumptions.

Our aim is to analyse the consequences of a change in the mathematical structure of characters on phylogenetic analysis, and its consequences for the resolution of coding issues raised by Pleijel. More precisely, we show that we can solve these issues using a hierarchical reformulation of phylogenetic hypotheses. To this end, we reformulate all coding approaches from Pleijel (1995) and from Forey & Kitching (2000) using hierarchies instead of partitions. We also propose new coding approaches, and produce a protocol through phylogenetic analyses in order to compare all of the approaches. We then show how the use of hierarchical characters allow workers to develop innovative types of coding that have no equivalent in partitions. Finally, we demonstrate how hierarchical characters allow to solve all the issues raised by Pleijel 25 years ago and that have never been solved since.

## Material and Methods

### Three-taxon analysis and hierarchical characters

In his original work, Pleijel defined characters and character states as “[…] *columns in data matrix in which taxa constitute rows, and character states the different values assigned to the entries*.” (Pleijel, 1995: 309). However, characters are more than columns, as he stated below: “*Information retrieval in this context relates to the efficiency of a coding procedure to transform observations without loss of data into a suitable form for phylogenetic analysis* […]” (Pleijel, 1995: 312), *i*.*e*. in characters. Characters are therefore intended to convey and order an information, the observations of taxa. We agree partially about this last point, but we argue that the concept of character is wider than columns in a data matrix. Columns are only a specific representation of the characters driven by a coding approach, but are not the characters themselves. Characters are hypotheses of relationships between taxa. Within this framework, character states are hypotheses of groupings of taxa based on homologs which are groupings of parts of taxa (Owen, 1848; Patterson, 1982). As characters are hypotheses on relationship between taxa, homologies are hypotheses on relationship between parts of taxa. Characters convey more than just strict observations because relationships are not observed in the nature, but are hypothesized on the grounds of observations. These hypotheses are constructed on the basis of the homologies. We will rely to the definition of characters as hierarchical hypotheses of relationships between taxa for the rest of this study.

Three-taxon analysis (3TA) is a method of phylogenetic reconstruction using hierarchical characters that lays in the cladistic paradigm. As for parsimony, 3TA uses characters to reconstruct a cladogram expressing phylogenetic relationships between taxa. This method is deeply rooted in a relational view of phylogenetic hypotheses. Clades and homologues characterise relationships between entities that are closer to each other than to any other. In their original paper, Nelson & Platnick (1991) proposed a new method to decompose character hypotheses in minimal kinship relationships between taxa that can be deduced from characters. These minimal phylogenetic units are sets of three taxa, named three-taxon statements (3TS), with two closers to each other than to a third, according to the character. The analysis in itself consists to analyse the total set of 3TS in order to reconstruct the cladogram (Nelson & Platnick, 1991; Rineau, 2017) in agreement with the maximum amount of 3TS. The congruence criterion is the maximum of compatibility between 3TS (equivalent to the minimum amount of rejected 3TS). The optimal tree is therefore the most informative about the relationship between taxa.

From this divergence, numerous methodological differences arise between 3TA and other phylogenetic methods such as standard parsimony. A difference between these two methods is the way standard parsimony and 3TA treat uncertainties, as polymorphism (Nelson & Ladiges, 1996; Zaragüeta Bagils et al., 2012), missing data and inapplicable data (Zaragüeta Bagils & Bourdon, 2007). In a parsimony analysis – using a classic partitional data matrix –, polymorphism, missing data and inapplicable data are all treated in the same way. Parsimony is not able to allow different treatments between these three types of uncertainty because of the underlying formalism of characters. Using a partition implies that every taxon necessary belongs to one set (*i*.*e*. the character state). If a taxon has no attributed state (because of inapplicable data or missing data), parsimony algorithms search to assign the taxon to the character state that consequently minimises the number of evolutionary steps of the most parsimonious tree(s). For missing data, the procedure violates the principle of character independence. Missing data issues are solved not by the worker, but by the algorithm, by means of a criterion (the number of steps of the most parcimonious tree(s)) which itself is related to the whole set of characters. The characters are then no longer independent (Zaragüeta Bagils & Bourdon, 2007). Although they are treated in the same way, the missing and inapplicable data do not contain the same information. A taxon with missing data means that we have no information about the particular trait: all states are possible. A taxon with an inapplicable data means that we have all the information to say that the taxon does not correspond to any state: no state is possible. These two opposite types of issues are confounded and treated the same way by parsimony algorithms. For polymorphism, we know that a taxon must be assigned to two or more states. By definition, partitions do not allow taxa to belong to multiple states. As a result, algorithms optimise in the same way as for missing and inapplicable data; we know that a taxon belong to several states but the procedure forces the taxon to fit only in one state.

However, hierarchies allow to distinguish the three different cases of uncertainties (missing data, inapplicable data, and polymorphism). i) We cannot hypothesise the relationship of taxa for whose we do not know their parts due to a missing data, because every state is possible. With the hierarchical representation, taxa with missing data are discarded from the character. The character tree will lack the corresponding leaves, which reduces the number of 3TS generated. All 3TS containing the discarded taxa are deleted so that the character does not contain any information on these taxa. As 3TS are units of information, a character with discarded taxa because of missing data will therefore be less informative than those that do not present this case. ii) When we face an inapplicable data for a specific taxon, the taxon is then connected to the root of the concerned character. The root of a hierarchical character does not represent a clear and defined state. It represents the speech universe of the analysis, based on the strong assumption of phylogenetic inference all taxa are linked by various degrees of kinship relationships. Therefore, we do not have to add an extra state of absence to speak about them: they are part of the analysis, but not included in this character. The analysis only reveals the different degrees of relationship between taxa, using shared character states. However, even an inapplicable case conveys information of these degrees: we know, for sure, the concerned taxa have not the feature. In other words, all other taxa are closer between them than the taxa with the inapplicable case, as they share the feature under its possible different forms. To connect taxa with inapplicable data to the root allow to generate 3TS between them and the other. Consequently, inapplicable data in a character will generate more 3TS than missing data, as they still allow to get some information where it is not possible for the latter. iii) Concerning polymorphism management using hierarchical characters, two different situations can be found. The first possibility is when one character state is included in another, *i*.*e*. when the first state is a particular case of the second. In this case, we can simplify by only considering the least inclusive character state because having this state is also possessing the more general one: in our example in fig 2a, *c* is present three times, *i*.*e*. it possesses three different states of the same character. Imagine now this character is about skin appendage and *c* possesses notably reptilian scales and feather. In this hypothesis, feathers are a particular kind of scales (the most derived *c* should be feather and one another *c* should be scales). Possessing feathers is then also possessing scales, under another state, more particular. Including both states does not teach us more than including only the more particular, as it is the most informative: it summarises the information from the other possibilities with its own information, which cannot be inferred in the opposite way to the above cases. We then keep 3TS including the more derived occurrence. In terms of 3TS, we will retain only 3TS from the more particular state. The other possibility is when several instances of the same taxa is present in mutually exclusive states (e.g. (ab)(ac)). There, one cannot be discarded, as both of them bring different information. In this kind of situation, we apply the paralogy free subtree analysis (Nelson & Ladiges, 1996). The character is split into several independent subtrees because mutually exclusive states are considered as independent, so there is no loss of information while cutting the tree into subtrees. Then, character subtrees avoid the repetition of taxa and allow to generate all the 3TS implied by the original case (fig 2b) (Cao, 2008; Morrone, 2009; Rineau, 2017).

**Figure 2.**
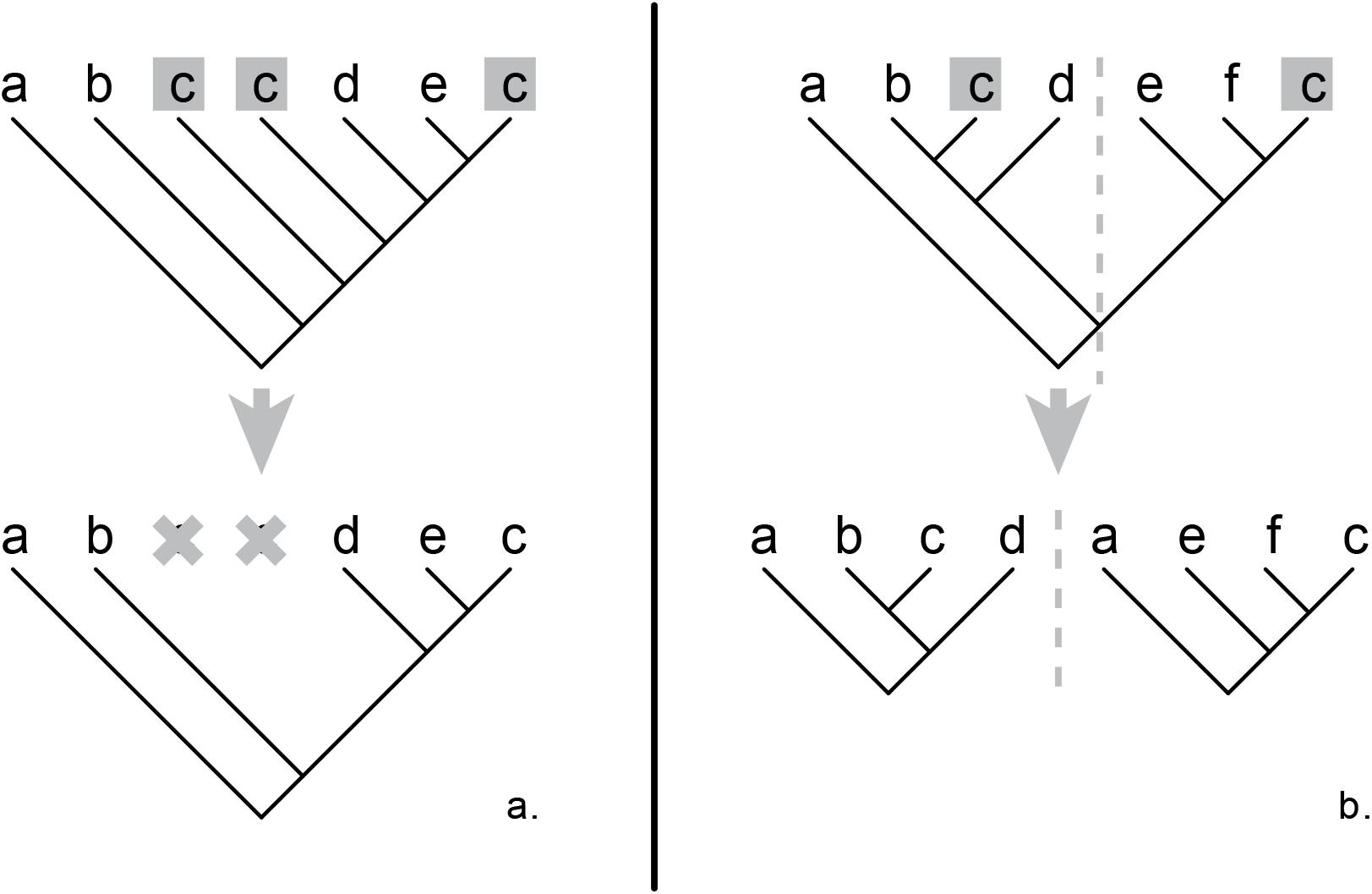
The two possible treatments of polymorphism in 3TA. In both case, c is present several times in the character because it has a polymorphic expression of it. In a., the repetition of c can be dealt with the simplification of the version of c branching close to the root. Keeping only the more derived c allows to keep the same information in term of i3TS. In b., we cannot proceed by the simplification of one or the other c, so, we have to keep both expression and treat them into two different subtrees. The separation of the subtree is made on the last common node for both c.

### Pleijel’s example

Pleijel’s (1995) feature X exists in four observable phenotypes: black and round, striped and round, black and squared and striped and squared. Pleijel stated also in his example that feature X is absent in some taxa (fig 1). In order to analyse the consequences of the different coding approaches on the reconstructed phylogenetic tree, we arose a total of ten taxa (named *a* to *j*), each phenotype being bore by two taxa: *c* and *d* have the black and round shape, *e* and *f* have the striped and round shape, *g* and *h* have the black and squared shape, and *i* and *j* have the striped and squared shape. The taxa *a* and *b* do not possess the feature X (fig 1). Using two taxa for each possibility allow each state to become synapomorphic, avoiding autapomorphies.

We transformed all proposed approaches of Pleijel (1995) and of Forey & Kitching (2000) into hierarchies, seeking to be as close as possible to their point of view. Furthermore, we propose here three new approaches that can only be coded in a hierarchical way. Neither Pleijel nor Forey & Kitching explicitly proposed a state as plesiomorphic. However, this assumption is necessary to ultimately discuss rooted trees, so we considered the absence of feature X as plesiomorphic in each case, as it is a common way to treat it. All coding approaches are detailed in fig 3 and 4 and in table 1. We use several criteria to compare the approaches. The number of independent three-taxon statements (i3TS) represents the amount of phylogenetic information an approach brings to the analysis, after a correction of its redundancy by fractional weighting. Each character being decomposed in 3TS produce redundant and overweighted statements that are not independent. The fractional weighting (Nelson & Ladiges, 1992; Williams & Humphries, 2003) is a method allowing to correct dependency issues by weighting 3TS. The i3TS therefore represents the amount of independent phylogenetic statements supported by an approach. It is the sum of 3TS corrected by fractional weighting of all characters of an approach. The number of optimal trees is another useful criterion to study the structure of the phylogenetic information conveyed by a coding approach. The less the number of optimal trees is, the more congruent and unambiguous the information is. Finally, the retention index (RI, Archie, 1989; Farris, 1989) adapted to hierarchical characters (Williams, 1998), is an efficient measure for assessing the rate of congruent information among trees. Originally, it was a measure of the homoplastic content of a character. In 3TA, the RI measures the rate of i3TS deduced from characters that are retained in the optimal tree:

**Table 1.**
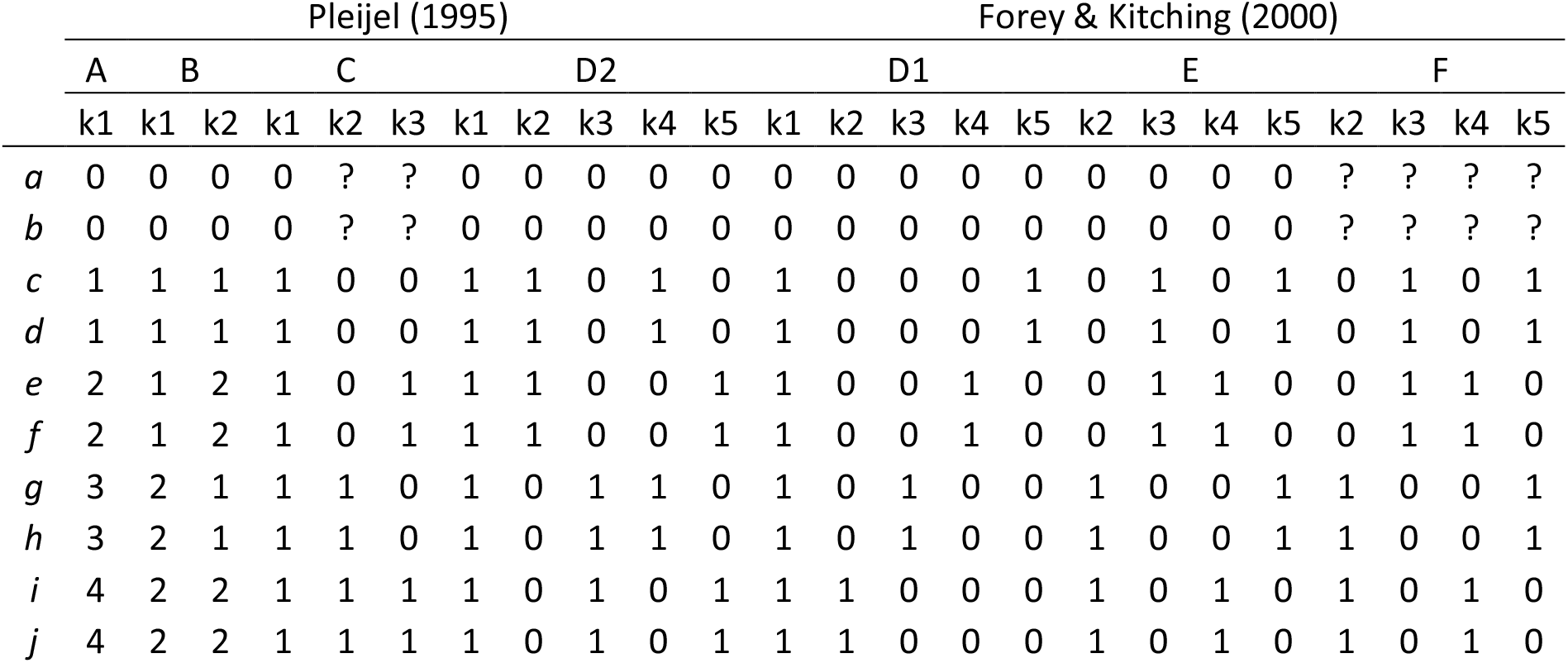
Pleijel’s and Forey & Kitching’s approaches in a matrixial version including our proposed taxa. A to F designate the different approaches with D1 and D2 sensu Forey & Kitching (D2 being Pleijel’s D approach). kn designates the different characters. 0 and 1 refer to the characters states and “?” to missing entries, as specified by both authors. (note that, for the rest of the paper, we will name Pleijel’s (1995) approach D as approach D2, sensu Forey & Kitching (2000))

**Figure 3.**
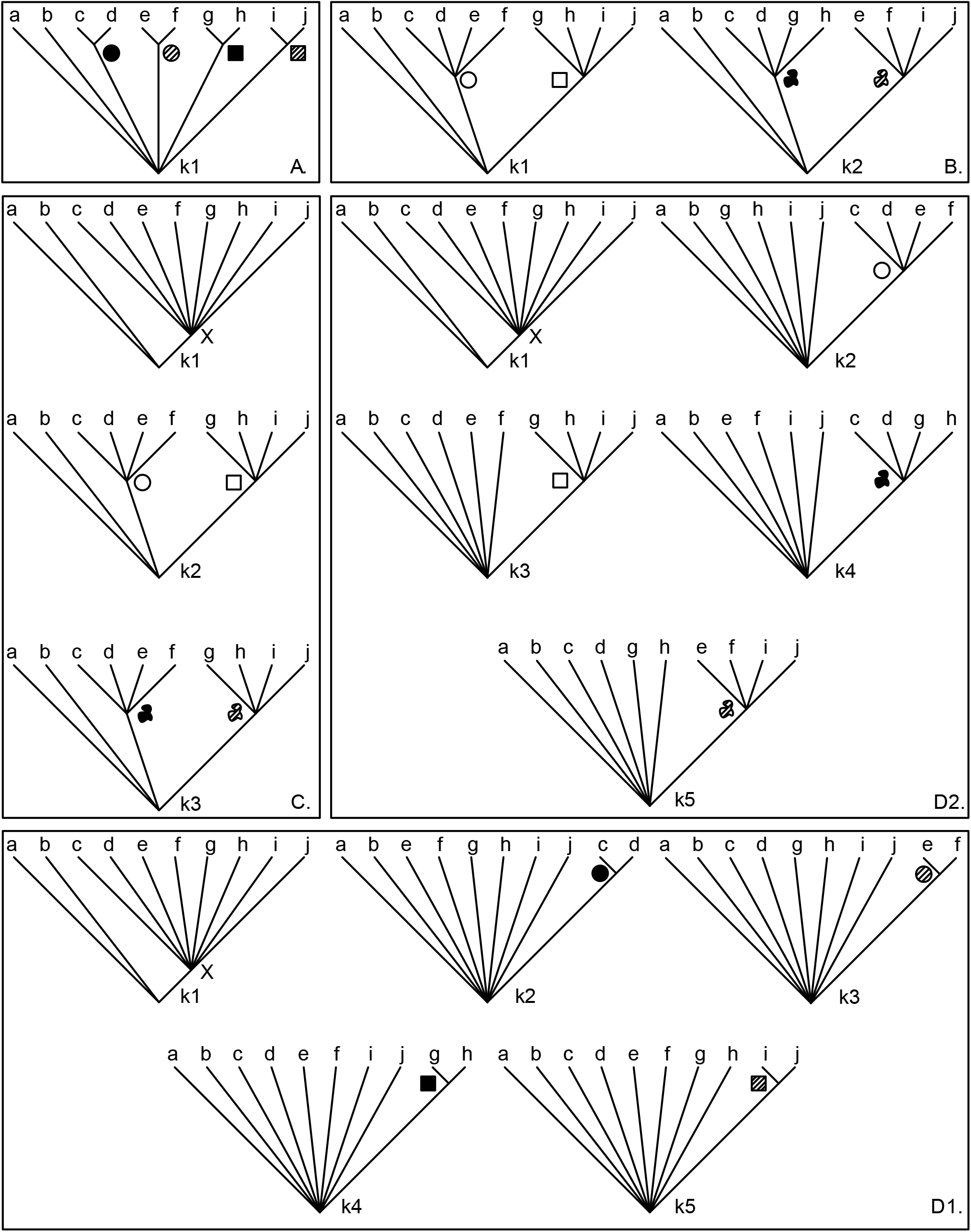
Translation of Pleijel’s A, B, C and D (under the name of D2) approaches, and of Forey & Kitching’s D1 approach. a to j are the taxa, kn are the different character contained in the approaches, and the figures designate the different states: X for the feature X itself, the black round, black square, striped round and striped square for the eponymous states, the empty square and round for the state square and round (respectively), and the black and striped spot-like figure for the state black and striped (respectively).

**Figure 4.**
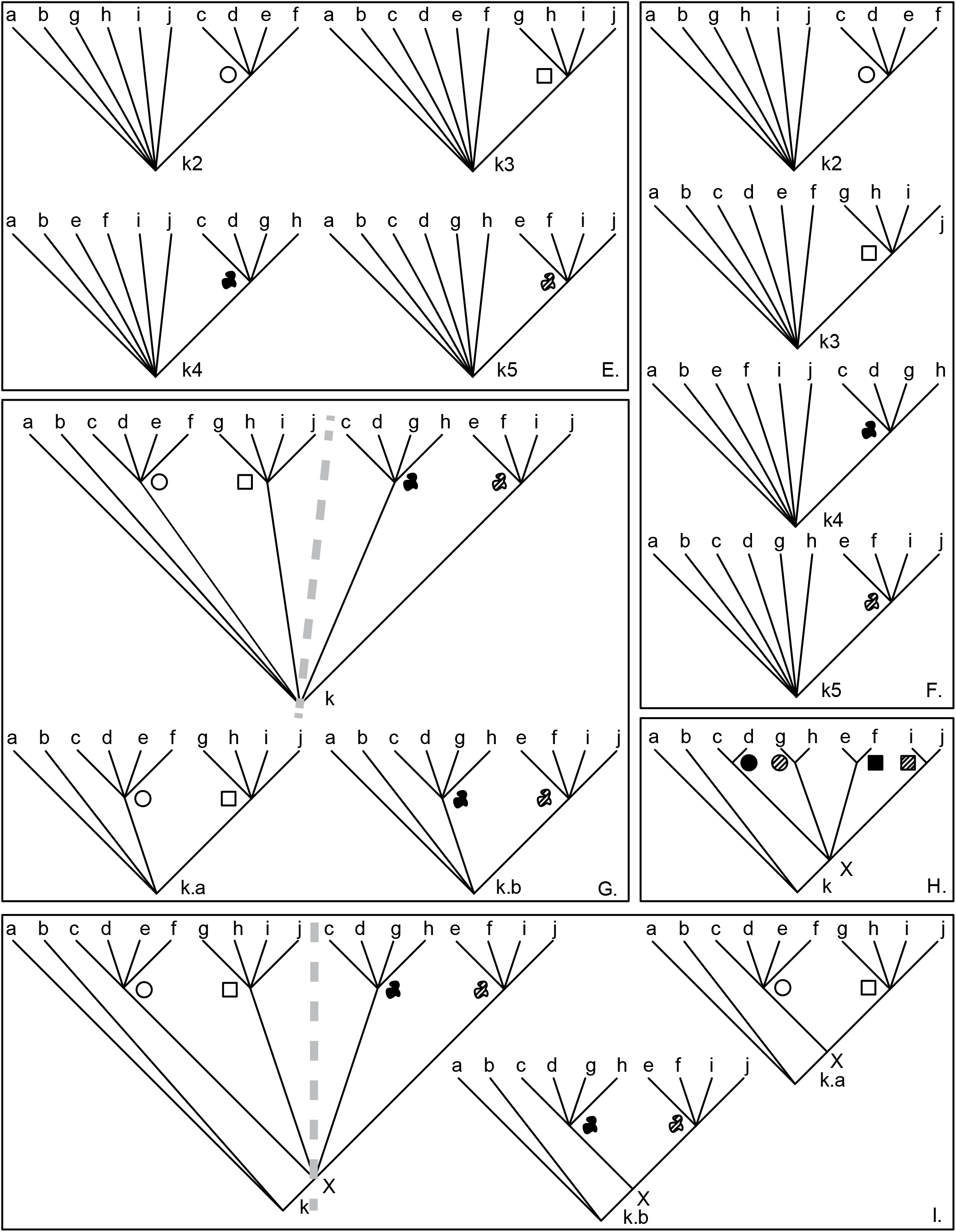
Translation of Forey & Kitching’s E and F approaches, and our own G, H and I approaches. a to j are the taxa, kn are the different character contained in the approaches (in case of G and I approach, k.a and k.b designate the subtree from k after free subtree paralogy), and the figures designate the different states: X for the feature X itself, the black round, black square, striped round and striped square for the eponymous states, the empty square and round for the state square and round (respectively), and the black and striped spot-like figure for the state black and striped (respectively).

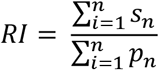

Where *n* is the number of characters, 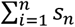 is the sum of the weight of all i3TS (*s*_*n*_) and 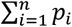 is the sum of the weight of all 3TS (*p*_*n*_). Incongruence appears when there are incongruent statements (*e*.*g. a*(*b,c*) and *b*(*a,c*)).

Less incongruent 3TS from one or several characters we have, closer to 1 will be the RI. On the opposite, a lower RI highlights a less congruent phylogenetic information. A high RI is to be preferred in order to avoid as many incongruences as possible. Contrary to i3TS, the RI score directly refer to the optimal trees from the used characters. The number of i3TS, the number of optimal trees, and the RI score were obtained through running phylogenetic analyses of each proposals with the software Lisbeth 1.0 (Zaragüeta Bagils et al., 2012). The paralogy free subtree analyses were directly performed using the ‘3area’ tool, included within the software. The RI score is measured on the different MPT, which share the same one when from the same analysis.

## Results

### Former approaches

We translated the characters proposed by Pleijel (1995) and then by Forey and Kitching (2000) into hierarchies. In binary character approaches, as in D1 and D2, the plesiomorphic state is the absence of the particular expression of feature X considered in the character. It is always true in multistate character approaches (*e*.*g*. B; k1: 0. absence, 1. black, 2. striped), but another question is raised here: how to ordinate the different states? In approach B, character 1, neither of states black nor striped includes the latter as a hierarchical subset. For example, black could be considered as a subset of striped, *i*.*e*. the latter being a derived state of the former. Although it is possible, we have no evidence (ontogenic, paleontological, etc.) to support such a speech. Therefore, these states are considered equals and none of them is included in the other. In approach C, question marks directly refer to an inapplicable state, as explained above: for k2 and k3, we linked taxa *a* and *b* directly to the root. Question marks of F are a non-applicable case, clearly stated by Forey & Kitching (2000: 58); it necessitates in hierarchical coding to branch the concerned taxa directly to the root. The approach E is a presence/absence coding with 0 meaning the absence of the character state. Therefore, E and F leads to identical hierarchical representations because they convey the same information, as represented in fig 4.

### New approaches

The first step for any worker in phylogenetic reconstruction based on morpho-anatomical data is to use comparative anatomy to propose homologs (groupings of parts), and then to deduce characters (groupings of taxa) based on it. Here, several groupings can be made:

- A grouping based on the sharing of the feature X, gathering all taxa from *c* to *i* (named “set X” hereinafter).

Then two possibilities of homologues recognition arise, leading to two options for grouping hypotheses. We can propose four grouping based on particular morphologies of X (set 1):

- rounded and black feature X (*c* and *d*)
- rounded and striped feature X (*e* and *f*)
- squared and black feature X (*g* and *h*) squared and striped feature X (*i* and *j*)

Other grouping can be made by considering the different pigmentations and shapes in isolation (set 2):

- rounded shape of feature X (*c, d, e* and *f*)
- squared shape of feature X (*g, h, i* and *j*)
- black pigmentation of feature X (*c, d, g* and *h*)
- stripped pigmentation of feature X (*e, f, i* and *j*)

Note that in any case taxa *a* and *b* directly rely to the root in a basal trichotomy because there is no information to resolve it. These last two sets of groupings support the same number of clades and, so, must be treated as equal. To discard one of them is to lose a potential phylogenetic information. Any approach supporting only one set of grouping, and not a combination of set X and one of the other sets, should be considered incomplete, and therefore potentially misleading.

There are only seven possible combinations of hypotheses to form a particular approach: i) set 1 only, ii) set 2 only, iii) set X only, iv) set 1 + set X, v) set 2 + set X, vi) set 1 + set 2 and vii) set 1 + set 2 + set X. Combinations i and ii are misleading because we would lose information without the inclusion of feature X. On the opposite, the combination iii is misleading because it does not take account of another set, giving more information. Moreover, we note here that coding particular associations of shape and pigmentation (set 1) and coding the different pigmentations and shapes as separate states (set 2) must be treated as two incompatibles sets of possibilities based on the same observations, excluding vi and vii. Combinations vi and vii are misleading as they lead to the coexistence of dependant and incompatible states. For example, the character state “rounded and black feature X (*c* and *d*)” would be logically dependent of “rounded shape of feature X (*c, d, e* and *f*)” and of “black pigmentation of feature X (*c, d, g* and *h*)” and would lead to redundancy biasing the analysis.

Three new approaches (G, H, I) are proposed here (fig 4) according to the hypothesised groups made above. We have no information on taxa about stratigraphic distribution, ontogenetic development, or any potential argument to prefer one particular combination as including another as a subset, *i*.*e*. a more derived state, in regards of the others, *e*.*g*., we have not enough information to propose a black feature to be a more derived form of a stripped one (or the reverse). So, our approaches represent the maximum subset the observation of taxa’s features alone allows. As these subsets gather taxa by the sharing of a particular character state, more the character contains subsets, more it will produce information for the phylogenetic reconstruction. Approach H and I differs from G by adding set X. In both G and H, free subtree paralogy is needed because of polymorphism. This polymorphism reflects the use of set 2 in the coding approach: it implies the presence of each taxa possessing the feature X two times, through their shapes and their pigmentations. Both expression of feature X, although differently distributed among taxa, are then polymorphic. We will show thereafter that in hierarchical coding, polymorphism is not necessary synonym of a misleaded coding and of a lower phylogenetic information content.

Anyway, it leads to incongruent i3TS: *e*.*g*., from H approach, we get (*i*(*c,e*)) from k.a and (*c*(*e,i*)) from k.b. It is not a particular case as we find incongruent i3TS for each taxon from *c* to *j*. They are already present in Pleijel’s and Forey & Kitching’s approaches, but separated in different characters.

### Metrics

The result of our analyses is summarized in table 2, and a detailed comparison of i3TS is joined in supp info. Among all approaches, the approach A has the worst number of i3TS (32) and of retained trees (945). Forey & Kitching’s (2000) D1 approach has a lower number of retained trees (45) and a slightly better i3TS score (46). All other approaches show only 6 optimal trees but have different i3TS scores. The B, E, F, and G approaches have 72 i3TS, H has 80 i3TS, C and D2 have 86 i3TS, and approach I reaches a maximal score of 100 i3TS. Both D1 and H have a maximal RI of 1. Approach I is the third most congruent approach (RI = 0.84) and B, E, F and G have the worst scores (RI = 0.778).

**Table 2.** Metrics values and the groups used in the approaches after the 3TA analyses. A to I designate the different approaches with D1 and D2 sensu Forey & Kitching (D2 being Pleijel’s D approach). i3TS is the amount of independent 3TS from all characters of the approach, number of trees is the number of optimal solutions after analyse of all characters of the approach, RI is the score measuring the proportion of i3TS kept in the optimal solutions and set is the sets used to construct the approach.

|  | Pleijel (1995) |  |  |  | Forey & Kitching (2000) |  |  | New approaches |  |  |
| --- | --- | --- | --- | --- | --- | --- | --- | --- | --- | --- |
|  | A | B | C | D2 | D1 | E | F | G | H | I |
| i3TS | 32 | 72 | 86 | 86 | 46 | 72 | 72 | 72 | 46 | 100 |
| Number of tree | 945 | 6 | 6 | 6 | 45 | 6 | 6 | 6 | 45 | 6 |
| RI | 1 | 0,556 | 0,628 | 0,628 | 1 | 0,556 | 0,556 | 0,556 | 1 | 0,68 |
| sets | 1 | 2 | X+2 | X+2 | X+1 | 2 | 2 | 2 | X+1 | X+2 |

## Discussion

### Information content of coding approaches

The different coding approaches, especially through their metric and the link with the sets used for their coding are described in table 2.

Approaches A, D1 and H have the poorest phylogenetic information content. Both have the lowest i3TS score, *i*.*e*. they allow to discuss very few relationships between taxa (32 for A, 46 for D1 and H). On the other hand, they reach the highest RI (RI = 1), which means that there is no incongruent information in the character. This apparent high consistency is correlated to the low amount of information of these approaches: less information obviously leads to less incongruence. Approaches A, D1 and H bring the fewest assumptions, as their clades gather taxa only two by two, using set 1 for A, and set 1 + set X for D1 and H. However, we learn nothing in the relationship between these groups of taxa and between the different shapes, as the consensus of the 945 optimal trees is a star phylogeny, or only with the node X in approaches D1 and H. The very high number of possible optimal trees is the direct cause of this large polytomy.

Approaches B, E, F and G show the lowest RI (RI = 0.778). They all use set 2 only in their coding, groupings by shape (squared and rounded) or pigmentation (black and striped). In other words, they all deal with partially incongruent character states because the different combination of shape and pigmentation are overlapping (*e*.*g*. any character state of shape and any character state of pigmentation have two taxa in common). Contrary to C, D2, and I, there is no character state allowing to support a clade by the presence of feature X in approach B, E, F and G (table 2), which makes these approaches incomplete. Obviously, adding the set X raises the number of i3TS. For example, D1 is equivalent to approach A plus set X, and consequently, D1 has more i3TS (46) and less possible optimal trees (45) than A (32 i3TS and 945 optimal trees). The same case is found for B, E, F and G, reaching 72 i3TS, while C and D2 both use the set 2 + set X and reach 86 i3TS. However, the number of optimal trees does not change between them (always at 6).

The preference for the set 1 coding strategy (A, D1, H) has the effect of greatly increasing the number of optimal trees, leading to a greater number of equiprobable scenario. The immediate effect is to confuse the result, generally a consensus, showing that no ambiguous scenario is possible. Excepted A, D1 and H, all approaches using the set 2 (with or without the feature X) reach the minimal number of six optimal trees. To include the set X does not necessarily lead to reach this minimum number of optimal trees, as we can see with D1 and H. The answer lies in the use of the set 2. Contrary to the set 1, the set 2 creates clades with a minimum of four taxa (contrary to the set 1 which creates clades of two taxa). Therefore, four clades of four taxa will necessarily create more i3TS than four clades of two taxa. Even if the set 2 leads to incongruent information, contrary of set 1 (all RI = 1 for A, D1 and H, and all other RI, of approaches using the set 2, are lower), this greater amount of information reduces the number of optimal solutions (6 against 45, even 945 for A). The goal of the systematist is to find the less possible optimal solution and then the more resolved consensus, we thus encourage the use of a set 2 for coding approaches. Given the number of taxa and our data, we cannot find less than six optimal trees. Taxa *a* and *b* cannot be included in any group gathering taxa *c* to *j* as there is no information supporting it. The algorithm finds three possible placements, solved in a polytomy branched at the root in the strict consensus. There is two incongruent solution for taxa *c* to *j*: either creating clades with the particular shape (squared or rounded) or creating clades with the particular pigmentation (black or stripped). As the incongruence between shape and pigmentation is unsolvable and placement of *a* and *b* will still be labile as we have no information using feature X, polytomies are unavoidable in the consensus given the starting parameters.

Not all solutions provide the same amount of information. With the exception of approaches A, D1 and H, the minimal number of i3TS generated is 72. From them, the approaches B, E, F, and G reach the lowest RI (0.778) and i3TS scores (72), meaning they bring a lot of incongruent information as they present contradictory assumption resulting in contradictory i3TS. This information come from their partially incongruent grouping of taxa. All of them use the set 2 approach but we showed clades grouped with shapes and pigmentation are contradictory. For example, in these approaches, referring to shapes, *c* is closer of *e* than *h*. However, referring to pigmentation, *c* is closer of *h* than *e*. It results in i3TS contradicting themselves. Moreover, none of these approaches imply the set X, contrary to C, D2 and I. If it does not reduce the number of optimal trees, including the set X is still important as it leads to a better RI score (RI = 0.814 for C and D2, 0.84 for I) and more i3TS (86 for C and D2, 100 for I). Approach I, then, maximises the number of i3TS (100 i3TS), and shows the best RI score (0.84) among the approaches leading to 6 optimal trees. In a wider analysis with more characters, this “extra” information would be relevant: the more i3TS the character gives, the more it will weight in the analysis, as i3TS directly rely with the fractional weighting. The number of i3TS should be then the main selection criterion for the best coding approach to use. The new I approach is close to the C / D2 approaches, by their use of set 2 and set X (table 2). In both cases, the new approach shows better metrics: it has more i3TS (100 against 86) and a better RI score (0.84 against 0.814). The only difference between old and new approaches is the full use of a nested hierarchy in the new one. The use of free subtree paralogy implies the creation of two subtrees, leading to the duplication of the node X. The presence of two nodes X increases the number of generated i3TS, correctly taking into account the fact that both paralogues support relationships. Without this duplication, approaches C, D2 and I are equals, by their use of both set 2 and set X. We encourage, then, the use of set 2 + set X coding approaches. If it brings incongruence, it also, paradoxically, brings more information, as all approaches using it produces more i3TS (from 72 i3TS to 100 for set 2 approaches; from 32 i3TS to 46 for set 1 approaches) and less optimal solutions (only 6 with set 2, at least 45 with set 1). Among approaches using set 2, those including set X brings more i3TS (72 with set 2 against 86 to 100 with set 2 + set X) and a better RI score (0.778 with set 2 against 0.814 to 0.84 with set 2 + set X). The best coding approach should, then, chosen among those using both set 2 and set X.

### Semantic content of coding approaches

The metrics used above make it possible to discuss comparisons between approaches on the base of quantitative data. In this part we will now discuss the different approaches from a qualitative point of view by analysing the semantic content of the different coding types. In particular, we will see if the dependency issues encountered by Pleijel still remains in hierarchical characters.

As enlighten by the different sets, the different morphologies of feature X can be treated in different ways: as an indivisible whole (set 1), as a particular expression of pigmentation and shape (set 2) and as the presence of the feature itself (set X). One major difference between the old (Pleijel’s and Forey & Kitching’s) and the new approaches is the way they treat the different set between them. In approach A, taxa are only gathered by the share of combinations of shape and pigmentation (set 1). It does not even gather these combinations as particular expressions of feature X (as B, E, F, G). One could argue this approach is precautionary. However, it does not bring a lot of information on the relationship between taxa. This approach only point the basic: the sharing of a particular morphology, as a black rounded feature X for example, but does not bring any information in the relationship between taxa possessing different morphologies of feature X. Adding a set X (as for D1) allow to discuss more relationship, by including the set 1 into the set X: set 1 states are just particular cases of the feature X itself, which seems pretty intuitive. A state is then a hypothesis of a clade and, as clades, multiple states can be included from the more general to the more particular. The same can be done with set 2 and set X, as in approach C or D2. A new problem arises: using a partition to code these approaches leads to create different characters discussing the same feature and violate the independence of characters (character linkage *sensu* (Pleijel, 1995)). This problem is especially true for the full binary approaches D1 and D2. As Pleijel stated, it solves many problems, especially hierarchical linkage. However, it leads to particular conceptual issues. In this logic, all particular shapes and pigmentations appear to be independent. In a matrix-based analysis, it leads to overweight the absence of the feature X, which will be represented in each character (Maddison, 1993; Platnick et al., 1991; Pleijel, 1995). Both D1 and D2 find an equivalent full hierarchical approach, in H and I respectively. In these approaches, the unification into one unique character allows to avoid the problem of independence between characters.

Our own approaches summarise all the coded information into one unique character as in approaches H and I which propose a unique character but with variations on the grouping strategy used (set 1 + set X or set 2 + set X). Whichever of these two approaches is chosen, they can solve the other issues raised by Pleijel. As we exemplified before, using hierarchical characters leads to avoid all the issues created by optimisations, as what Pleijel called missing entries, *i*.*e*. the fact a “?” does not allow to distinguish the cases of polymorphism, missing data and inapplicable data. As there is only one character, all issues linked to “hierarchical linkage”, *i*.*e*. the different ways of coding the same feature in different characters, (*sensu* Pleijel, 1995: 310-311) vanish. Finally, as shown by our metrics, the information retrieval and testability, *i*.*e*. the efficiency to code phylogenetic information, is maximal when a hierarchical representation is used. This last point is especially dependent of the use of set 1 or 2, with or without a set X, as shown with our metrics.

Then, using a unique hierarchal character instead of several partitional ones (like I against C) fulfils all Pleijel’s criteria. Moreover, we can then propose one approach among the other as the best to use. As detailed before, one should choose a hierarchical approach using both set 2 and set X coding. This kind of approach is the answer sought by Pleijel in 1995.

## Conclusion

Prin (2012) proposed a reinterpretation of the cladistic theory focusing on clades as the product of evolution. Under this paradigm, only one tree (a unique inclusive hierarchy of clades) is the epistemic access phylogenetics allows to the course of evolution. The cladistic analysis is, so, a methodology aiming to find this tree. As every analysis, it proceeds by decomposing the problem into several simpler question, find the solutions to these questions, and synthesising all partial solution in the complete one, answering to the starting one (Cao et al., 2007; Descartes, 1637; Zaragüeta Bagils & Bourdon, 2007). However, no information could be added after the proposition of the partial solutions. In other words, all the information in the global solution is already contained in the partial ones. The global solution is no more than the sum of the partial ones. In cladistic analysis, according to Prin (2012) and Rineau (2017), the partial solutions are the characters, synthetized through the algorithm into the global solution, the cladogram. A problem arises: if the final tree is hierarchical, the partial solutions, the characters, should be either (Bock, 1963; Patterson, 1982). If not, it will be adding information and violating the principle of the analysis. Under this point of view, partitional characters could not be considered as cladistics and should be discarded for the benefit of hierarchical ones. States of character are clade hypotheses and are naturally represented as hierarchies. Therefore, if a character is a phylogenetic hypothesis, then a hypothesis of phylogeny, an optimal solution, is naturally a hierarchical tree.

Hierarchies present numerous advantages on partitions in a cladistic analysis. They make it possible to deal appropriately with the various problems of uncertainty (missing data, inapplicable data and polymorphism), where partitions could not solve them. Hierarchical characters also have the undoubted advantage of being able to be analysed thanks to the i3TS, which allows to quantify their phylogenetic information content, and to the RI, which allows to quantify their congruence, by measuring the part of i3TS retained in the optimal solution. These different tools allow both to apprehend the semantic content of the cladistic hypotheses, and to allow the worker to choose efficiently and precisely between several codings. To use biogeographic methods, such as 3TA (Nelson & Ladiges, 1992, 1996; Nelson & Platnick, 1991; Zaragüeta Bagils et al., 2012), to solve this problem made it understandable and offered a solution for the first time. It is the better way to represent character, as it solves all Pleijel’s (1995) issues and as states of characters, as hypotheses of clades, are naturally hierarchical.

## Acknowledgements

We would like to thank CR2P for the funding that made this study possible. We also would like to thank Stéphane Prin, Jérémie Bardin, Bouziane Khalloufi, Artémis Korniliou, Paul Zaharias and Paul Chatelain for fruitful conversation on these topics. We thank Jorge Cubo for his help and comments about this manuscript. Finally, we thank Pascal Tassy for fruitful comments on this work. R.Z.B and V.R. designed this study, M.G.F.-B. conducts the experiments and wrote the first manuscript. All three authors contributed equally to the final version of this paper.

